# Glutamatergic systems in Hydrozoa (Cnidaria)

**DOI:** 10.64898/2026.08.23.746573

**Authors:** Leonid L. Moroz, Tigran P. Norekian

**Affiliations:** Whitney Laboratory for Marine Bioscience, University of Florida, St. Augustine, FL, 32080, USA; Department of Neuroscience and McKnight Brain Institute, College of Medicine, University of Florida, Gainesville, FL 32611

**Author notes:** **Corresponding author**: Leonid L. Moroz.

**Keywords:** Cnidaria, Siphonophores, Ctenophora, Nervous system evolution, Nerve nets, Nematocytes, Evolution of neurotransmitters, glutamate and GABA signaling, muscle physiology, *Aglantha*, *Polyorchis*, *Aequorea*, *Clytia*, *Nanomia*

## Abstract

The origins and early diversification of intercellular signaling molecules in animals remain poorly understood because comparative data across basal metazoan lineages are limited. Cnidarians form the sister group to bilaterian animals, and characterizing their transmitter systems is critical to understanding how complex adaptations within integrative systems shape evolutionary trajectories. Although glutamate is a well-established transmitter in bilaterian animals, its role in cnidarians remains unclear, and information on its neuronal function and signaling is limited. For most studied cnidarians, glutamate has been suggested to be a non-neuronal signaling molecule. Here, using glutamate immunoreactivity (Glu IR) in eight hydrozoan species with distinct ecologies (*Aequorea victoria, Eutonina indicans, Clytia gregaria, Bougainvillia principis, Euphysa flammea, Polyorchis penicillatus, Aglantha digitalis, Nanomia septata*), we identified and visualized distinct populations of glutamate-immunoreactive (Glu-ir) cells, including nematocytes, neurons, and muscle cells. A broad diversity of Glu-ir nematocytes was found in all studied species. Glu-ir neural cells were found only in three species (*Aequorea, Nanomia*, and *Aglantha*); their morphology and localization were species-specific. In addition, some striated and smooth myoepithelial cells were found to be either Glu-ir or GABA-ir. We propose that both glutamatergic and GABAergic systems were independently recruited more than 3 times as neurotransmitters across cnidarians, and that these recruitments are fundamentally rooted in bioenergetic demands.

## INTRODUCTION

The origins and early diversification of intercellular signaling molecules remain poorly understood because comparative data across basal metazoan lineages are limited (Moroz, 2014, 2021). Cnidaria is the sister group to bilaterian animals and is critical for understanding both neuro-ecological adaptations and the dynamics of integrative systems along different evolutionary trajectories. Historically, studies of cnidarians have provided the first insights into the evolution of neural systems (Anctil, 2015), from early descriptions of neural nets to the identification of multiple parallel conductive systems for behavioral integration (Mackie, 1970, 1990) and molecular architectures (Kelava et al., 2015; Bosch et al., 2017). However, chemical transmission has received less attention because of numerous analytical challenges and the enormous diversity, especially among hundreds of neuropeptides. About a dozen small-molecule transmitters are known in bilaterians; however, data for cnidarians are often controversial (Anctil, 2009).

Comparative genomics suggests that cnidarians lack canonical transmitters such as serotonin, dopamine, noradrenaline, adrenaline, histamine, and acetylcholine (Moroz, Kohn, 2015; Moroz et al., 2021a), although other endogenous monoamines may exist. The gaseous messenger nitric oxide (NO), glutamate, and gamma-aminobutyric acid (GABA) are claimed to be operational in a diverse range of cnidarians (Anctil, 2009), but their neuronal localization is contested. For example, the neural location of NO synthase and its function were confirmed in only one species among many studied (Moroz et al., 2004). The neuronal localization of GABA (Marlow et al., 2009) and glutamate (Delgado et al., 2010) was reported for two species of Anthozoa, but non-neuronal localization and function of these messengers appear to be predominant features in cnidarians (Carlyle, 1974, Anctil and Carette, 1994; Oren et al., 2014).

This situation presents an opportunity to reconstruct transmitter recruitment from non-neuronal exaptations, under the hypothesis that transmitter origins are rooted in core metabolism. We focused primarily on glutamate and GABA: metabolically coupled anaplerotic molecules that refuel the Krebs cycle (tricarboxylic acid cycle or TCA), the universal bioenergetic architecture for life. Glutamate is the most abundant intercellular metabolite, whereas GABA is a product of glutamate decarboxylation -- glutamate decarboxylase (GAD) catalyzes the irreversible synthesis of GABA (Andersen, 2025).

Glutamate was hypothesized to be one of the most ancient signaling molecules, linking metabolism, excitability, and intercellular communication, from prokaryotes to eukaryotes (Moroz et al., 2021b). In the context of neural evolution, glutamate was recruited as the neuromuscular transmitter in ctenophores (Moroz et al., 2014); it was localized in the subepithelial nerve net and in some mesogleal neuroid cells (Moroz, Norekian, 2026). In contrast, neural localization of GABA was not demonstrated in ctenophores. GABA was also ineffective in modulating ciliary beating in semi-intact preparations (Norekian and Moroz, 2024) and did not induce contractions of muscles sensitive to glutamate (Moroz et al., 2014). In other words, the neurotransmitter function of GABA is not (yet) supported in ctenophores.

The main question of this study is whether glutamatergic and GABAergic neurons exist in hydrozoan medusae. Here, we used glutamate- and GABA-specific antibodies to characterize the distribution of two putative intercellular messengers in eight pelagic species of the class Hydrozoa, representing both individual medusae and colonies. Although the phylogeny of Hydrozoa is not fully resolved, the selected species illustrate substantial diversification of swimming and feeding adaptations, with distinct ecologies and locomotory behaviors. The observed mosaic, cell-specific recruitment of glutamate and GABA across neural and muscular systems reflects a highly dynamic interplay of energetic and signaling constraints in the evolution of these intercellular transmitters beyond traditional synaptic communication. We propose that glutamatergic neurons evolved at least four times independently within Cnidaria.

## MATERIALS and METHODS

### Animals

Adult specimens (Fig. 1) of seven hydrozoan jellyfish species and a colonial siphonophore (Fig. 4A) were collected from the breakwater in the Northwest Pacific Ocean at Friday Harbor Laboratories, University of Washington: *Aequorea victoria* (Leptothecata, Aequoreidae), *Eutonina indicans* (Leptothecata, Eirenidae), *Clytia gregaria*(Leptothecata, Campanulariidae), *Bougainvillia multitentaculata* (Anthoathecata, Bougainvilliidae), *Euphysa flammea* (Anthoathecata, Corymorphidae), *Polyorchis penicillatus* (Anthoathecata, Corynidae), *Aglantha digitalis* (Trachymedusae, Rhopalonematidae), and *Nanomia septata* (Siphonophorae, Agalmatidae). Specimens were kept in 1-gallon jars at 10 °C for no more than 4-5 days before processing.

**Figure 1.**
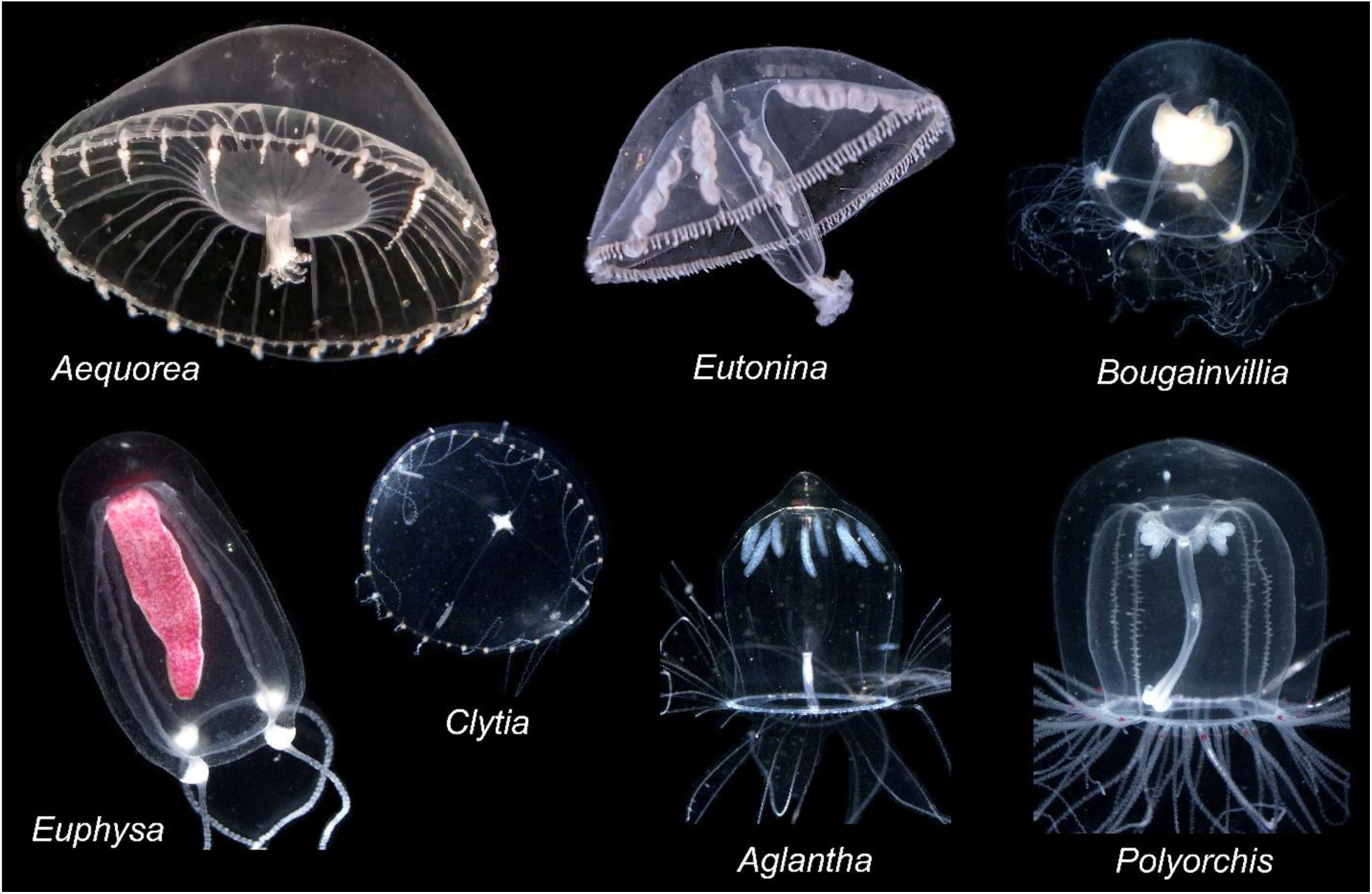
Hydrozoan medusae used in the current study (Class Hydrozoa). *Aequorea victoria* (Leptothecata, Aequoreidae)*, Eutonina indicans* (Leptothecata, Eirenidae)*, Clytia gregaria (Leptothecata, Campanulariidae), Bougainvillia multitentaculata* (Anthoathecata, Bougainvilliidae)*, Euphysa flammea* (Anthoathecata, Corymorphidae), *Aglantha digitalis* (Trachymedusae, Rhopalonematidae), *Polyorchis penicillatus* (Anthoathecata, Corynidae). These hydrozoan medusae are common in the Northwestern Pacific.

### Immunocytochemistry and Phalloidin staining

Adult animals were fixed for 10-14 hours in 4% paraformaldehyde in 0.1 M phosphate-buffered saline (PBS) at +5 °C. They were then washed for 4-6 hours in PBS and dissected into smaller pieces. Fixed tissues were pre-incubated in a blocking solution of 6% goat serum in PBS containing 0.02% Triton X-100 (PBT) for 12 hours. Samples were incubated for 48 hours at +5 °C with primary antibodies diluted in 6% goat serum to a final dilution of 1:200. We used polyclonal anti-glutamate antibodies produced in rabbit (SigmaAldrich, Cat#G6642, RRID: AB_259946). Following a series of PBS washes for 6-8 hours, tissues from adult animals were incubated for 20 hours in secondary goat anti-rabbit IgG antibodies conjugated to Alexa Fluor 568 (ThermoFisher Cat# A-11011, RRID: AB_143157) at a final dilution of 1:100. To identify GABA immunoreactivity, we used polyclonal anti-GABA antibodies produced in rabbit (SigmaAldrich, Cat# A2052, RRID: RRID: AB_477652). The same secondary goat anti-rabbit antibodies were used for the GABA primary antibodies.

We also used rat monoclonal antibodies against tubulin (AbD Serotec Cat# MCA77G, RRID: AB_325003) in double-labeling experiments at a final dilution of 1:100 to identify overall neuronal populations. These antibodies recognize the alpha subunit of tubulin and specifically bind tyrosylated tubulin (Wehland & Willingham, 1983; Wehland et al., 1983). After a series of PBS washes lasting 6-8 hours, the dissected tissues were incubated for 12 hours with secondary goat anti-rat IgG antibodies conjugated to Alexa Fluor 488 (Molecular Probes, Invitrogen, Cat# A11006, RRID: AB_141373) at a final dilution of 1:100.

To label the muscle fibers, we used the well-known marker phalloidin (Alexa Fluor 488 phalloidin from Molecular Probes), which binds to F-actin. After washing in PBS following secondary antibody treatment, the samples were incubated in phalloidin solution (in PBS) for 8 hours at +5°C at a final dilution of 1:100, then washed with several PBS rinses for 6-8 hours. then washed in several PBS rinses for 6-8 hours.

To stain the nuclei, dissected tissues were mounted in Antifade Mounting Medium containing DAPI (Vectashield; Cat#H-2000). Slides were examined on a Nikon Research Microscope Eclipse E800 with epifluorescence using standard TRITC and FITC filters, and images were recorded on a Nikon C1 laser-scanning confocal microscope.

## RESULTS

The most prominent cell-specific glutamate IR labeling was observed in nematocysts (stinging cells) across all studied cnidarians, regardless of their localization and morphology (**Figs. 2, 3, 5**). Nematocysts are the hallmark of the phylum Cnidaria, and these cells have previously been recognized for their high glutamate content (Weber, 1990).

**Figure 2.**
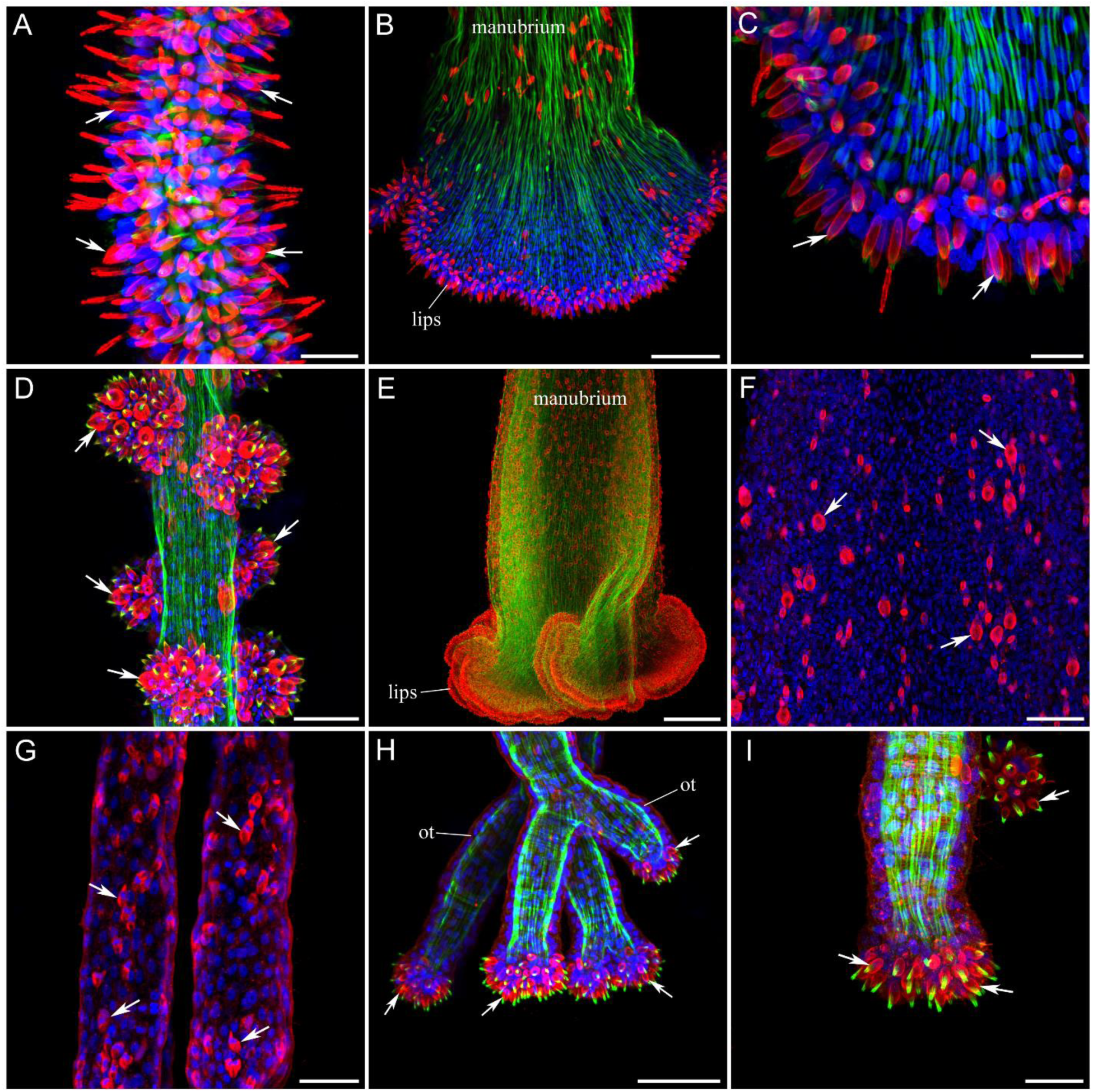
Nematocysts in Hydrozoans. Nematocysts are brightly labeled with Glutamate IR (red) in *Eutonina* (A, B, C), in *Polyorchis* (D, E, F), and in *Bougainvillia* (G, H, I). Phalloidin labeling is green, while DAPI is blue. A – *Eutonina’s* tentacle is densely covered with nematocysts (arrows). B – The manubrium, and especially the lip area, contains many nematocysts labeled by Glu IR in red. C – Higher magnification of the lip area. Some of the nematocysts are indicated by arrows. D – Nematocysts (arrows) are clustered in large groups along the *Polyorchis* tentacle. E – Manubrium and lip area. F – Higher magnification of the manubrium surface reveals numerous nematocysts (arrows) labeled with Glutamate IR. G – Tentacles in *Bougainvillia* have numerous nematocysts (arrows) spread along their entire length. H, I - Unique to *Bougainvillia* oral tentacles have large clusters of nematocysts (arrows) at the end of each tentacle. Scale bars: A – 15 µm, B – 50 µm, C – 10 µm, D – 50 µm, E – 200 µm, F – 50 µm, G – 30 µm, H – 50 µm, I – 20 µm.

**Figure 3.**
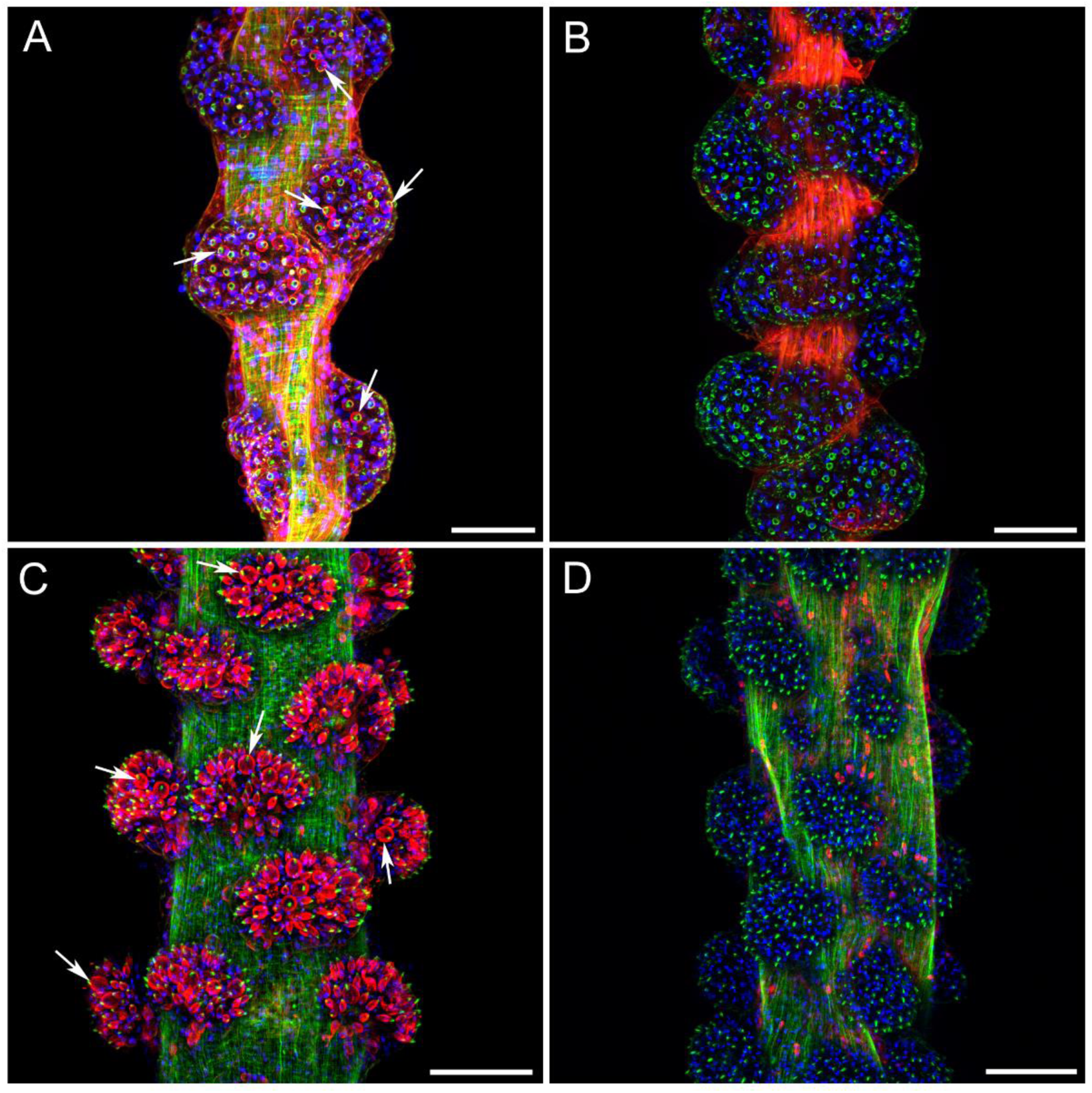
Glu and GABA immunoreactivity in the tentacles. Nematocysts in the tentacles of *Euphysa* (A, B) and *Polyorchis* (C, D). Glutamate antibodies label nematocysts (arrows) in both *Euphysa* (A) and *Polyorchis* (C), while GABA antibodies do not label nematocysts (B, D), but label other cell types such as muscles in *Euphysa* and neuron-like cells in *Polyorchis*. Glutamate IR and GABA IR are red, while phalloidin is green (DAPI is blue). Scale bars: 100 µm.

**Figure 4.**
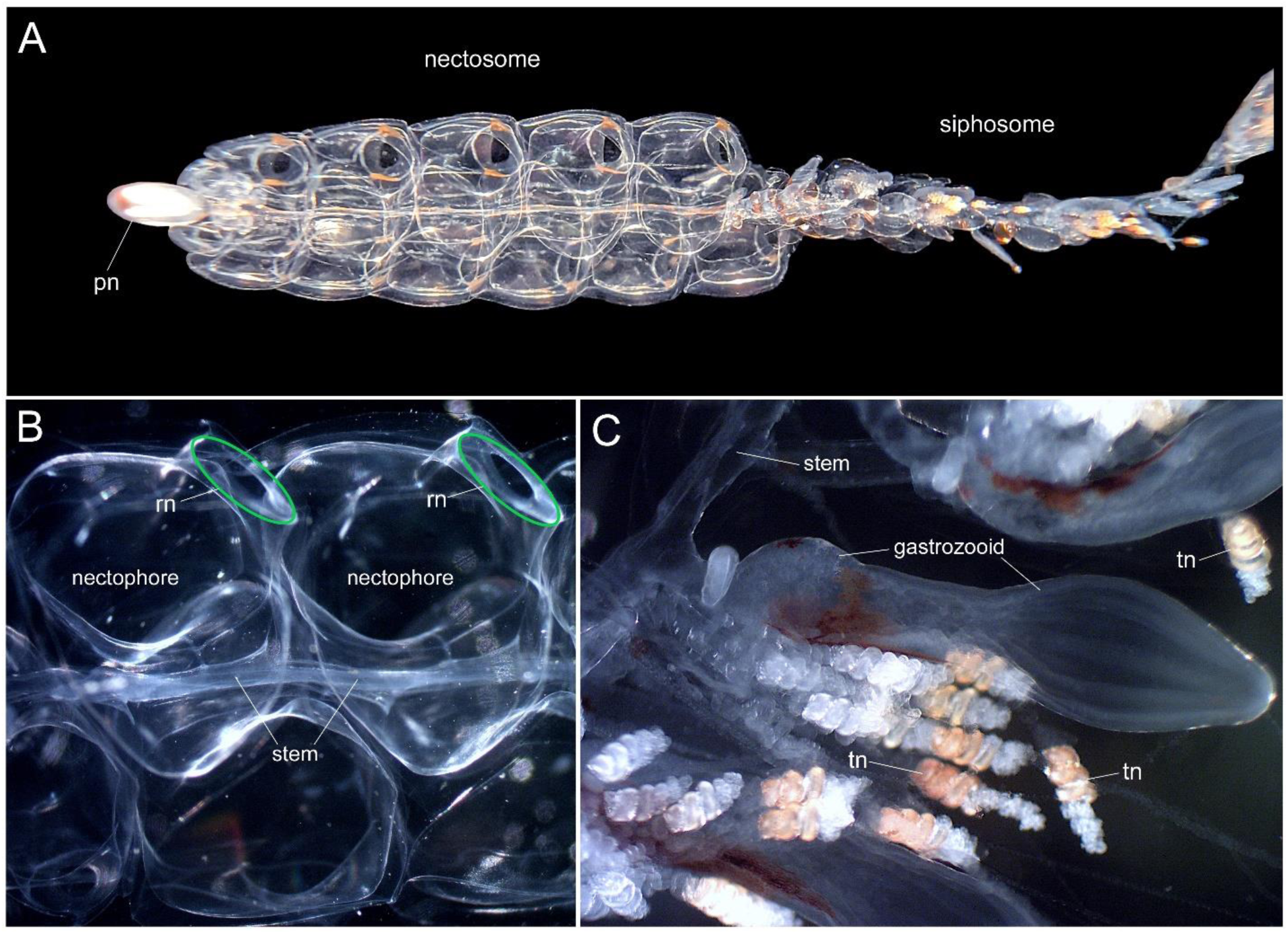
Live *Nanomia septata* and two major zooid classes – nectophores and gastrozooids. A – Entire *Nanomia* colony consists of two main regions: nectosome, which includes all nectophores (swimming bells) and siphosome with all other zooids. Pneumatophore (pn) is at the anterior end of the colony. B – Higher magnification of two nectophores with the location of the ring nerve (rn) shown schematically in green. C – Each gastrozooid in the siphosome area includes, in addition to the main body, a long tentacle with multiple branching tentilla (tn) used to capture prey.

**Figure 5.**
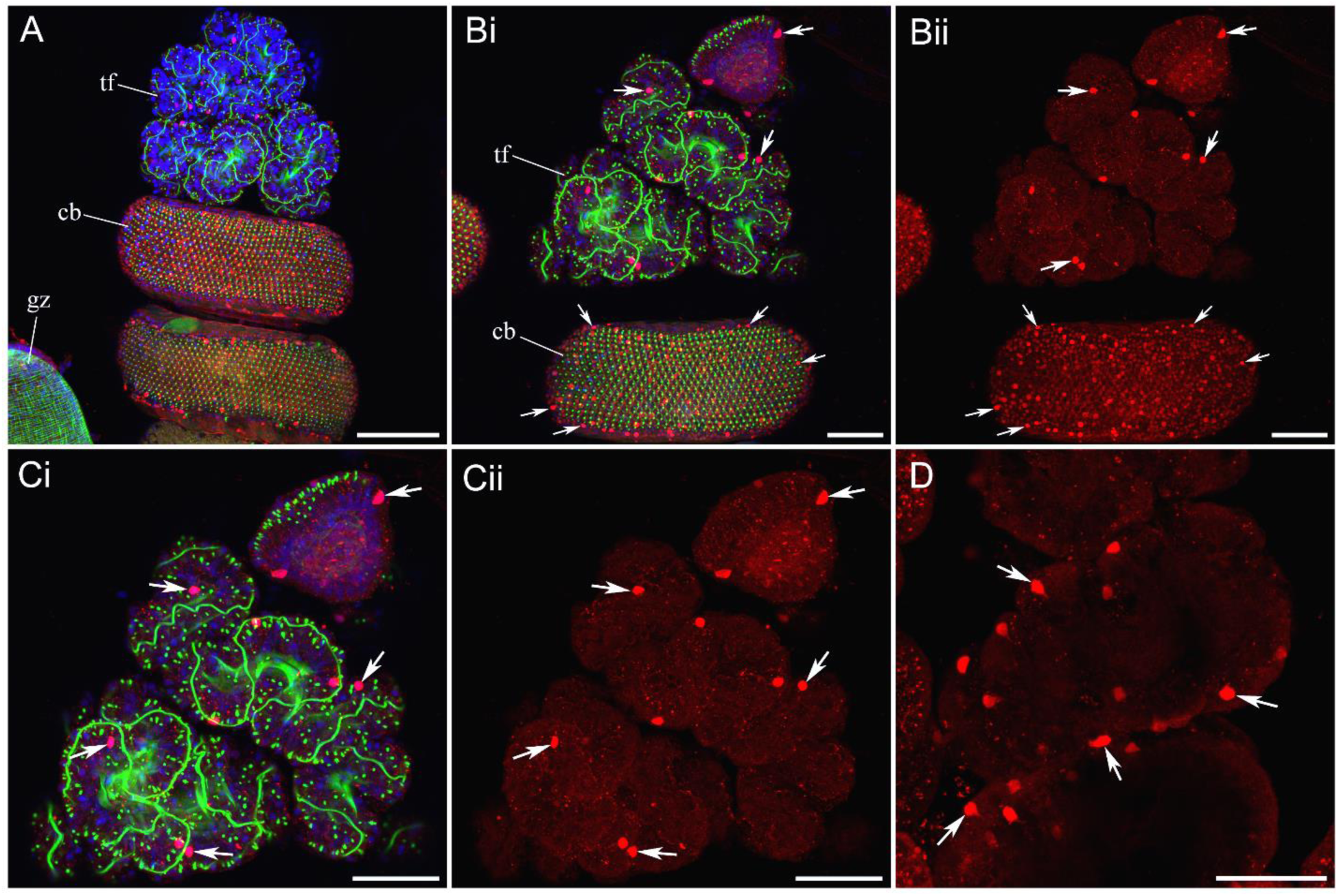
Glutamate IR in *Nanomia* tentacles. Glutamate immunoreactivity in the *Nanomia* tentilla. Glutamate IR is red, phalloidin is green, and DAPI is blue. A – Individual mature tentillum, which is a part of a gastrozooid (gz), includes a spiraled cnidoband (cb) with a battery of nematocysts and a terminal filament (tf). B - Numerous nematocytes on the outer surface of the spiraled cnidoband (cb) are labeled with glutamate AB in red (some are indicated by small arrows). In addition, larger spherical Glutamate-ir cells (arrows) are found in the terminal filament (tf). Bii - Red channel of Bi with Glutamate IR only. C - Higher magnification of the terminal filament shows the Glutamate-ir cells (arrows). Cii - Red channel of Ci with Glutamate IR only. D – Some of these Glutamate-ir cells in the terminal filament have a process branching from their cell body. Scale bars: A – 100 µm, B – 50 µm, C – 50 µm, D – 50 µm.

Nematocysts were found in the tentacles and in the manubrium around the mouth. In *Eutonina* (**Fig. 2A**) and *Bougainvillia* (**Fig. 2G**), Glu-ir nematocysts were evenly distributed along the entire length of the tentacles. In *Polyorchis* (**Fig. 2D; 3C**) and *Euphysa* (**Fig. 3A**), nematocysts were grouped into spherical clusters on the tentacles, a characteristic feature of these medusae. In the colonial siphonophore *Nanomia*, tentacles with branching tentilla are attached to gastrozooids, the feeding zooids (**Fig. 4C**). Each tentilla has a spiral cnidoband with a battery of nematocysts, which were also labeled with Glutamate IR (**Fig. 5A, B**). Glu-ir nematocysts were also found in the manubrium of all species studied. They had the highest density in the lip area around the mouth but were also present along the entire length of the manubrium – *Eutonina* (**Fig. 2B, C**), *Polyorchis* (**Fig. 2E, F**). Unlike other species, *Bougainvillia* has a set of unique oral tentacles with Glu-ir nematocysts concentrated in dense clusters at the very end of each tentacle (**Fig. 2H, I**). In all instances, glutamate IR showed a very strong signal from the cell bodies of all nematocysts studied, suggesting a high Glu concentration in these cells. In contrast to glutamate IR, GABA IR didn’t label nematocysts (**Fig. 3B, D**), presumably indicating the absence of GABA in nematocysts at concentrations sufficient for immunohistochemical detection. GABA IR experiments also served as a good control for glutamate IR, since the secondary antibodies and the entire protocol were the same as for glutamate IR.

Nematocytes and neurons share a common early developmental lineage as two distinct secretory cell types (Babonis et al., 2022; Steger et al., 2022). However, only a small subset of neurons was found to be glutamatergic. We observed glutamate IR in only three species studied (**Figs. 5-9**). The most prominent and intense glutamate IR was detected in *Nanomia* nectophores, specialized swimming zooids responsible for jet propulsion of the siphonophore colony (**Fig. 4 B**). These glutamate-ir neurons were located next to the ring nerve and appeared to be part of this ring neural system that controls the generation and modulation of the swim rhythm (**Fig. 6A**). The Glu-ir ring neurons had clearly identifiable processes; some were unipolar, while others were bipolar or multipolar, with branching processes and complex morphology (**Fig. 6 B, C**). In *Aequorea*, glutamate immunoreactivity was found in the subumbrella nerve net, staining a few scattered usually bipolar or tripolar neurons with sensory-like morphology – many of them had a short apical transepithelial process (**Fig. 7**). The density of these neurons was low, and most of the network was not Glu-ir. Finally, in the trachymedusa *Aglantha*, we identified two distinct neuronal subpopulations. One subpopulation was distributed along the length of the tentacles, with predominantly bipolar and some multipolar neurons (**Fig. 8**). The second subpopulation was identified at the base of *Aglantha*’s tentacles, forming two rows of sensory cells running longitudinally (**Fig. 9A, B**). These cells have been previously identified in *Aglantha* as sensory neurons, each projecting a long sensory cilium (see Norekian and Moroz, 2020). We have now learned that they are also glutamate-ir.

**Figure 6.**
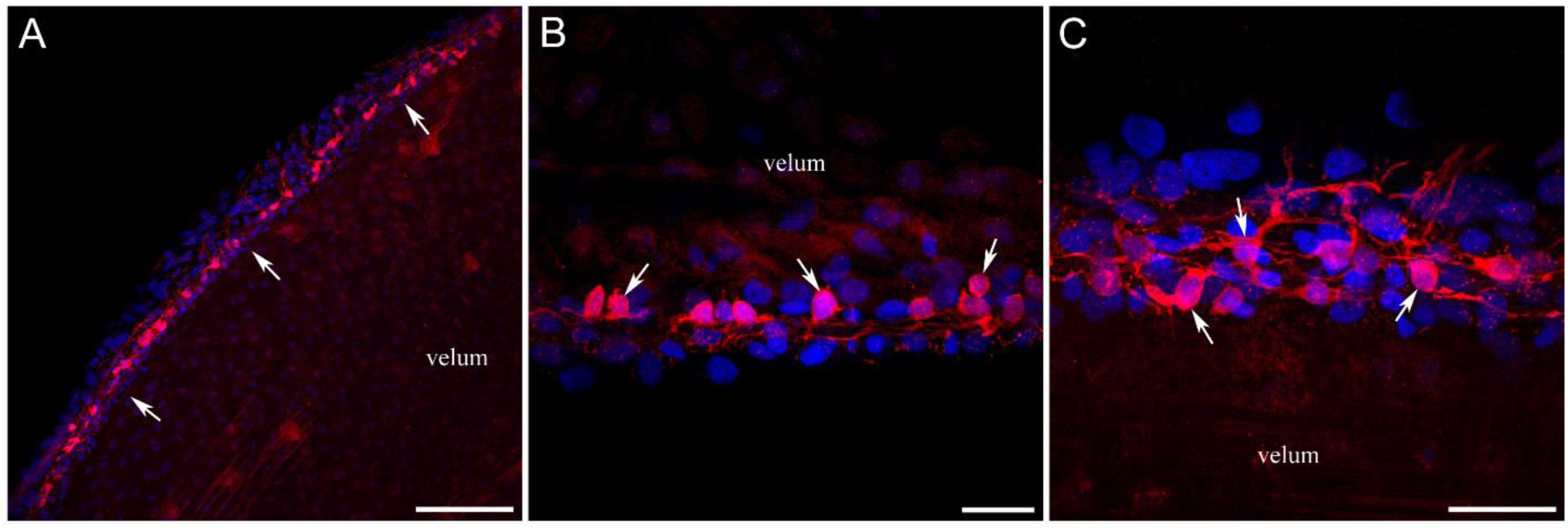
Glutamate-ir neurons around the ring nerve in *Nanomia* nectophores. Glutamate-ir neurons around the nerve ring in *Nanomia* nectophores. A – Immunoreactive neurons along the nerve ring (indicated by arrows) at the edge of the velum. B – Glutamate-ir neurons (arrows) at the lower section of the nectophore. C – Glutamate-ir neurons (arrows) at the upper section of the nectophore. Note the more extensive neural branching. Scale bars: A – 100 µm, B – 25 µm, C – 25 µm.

**Figure 7.**
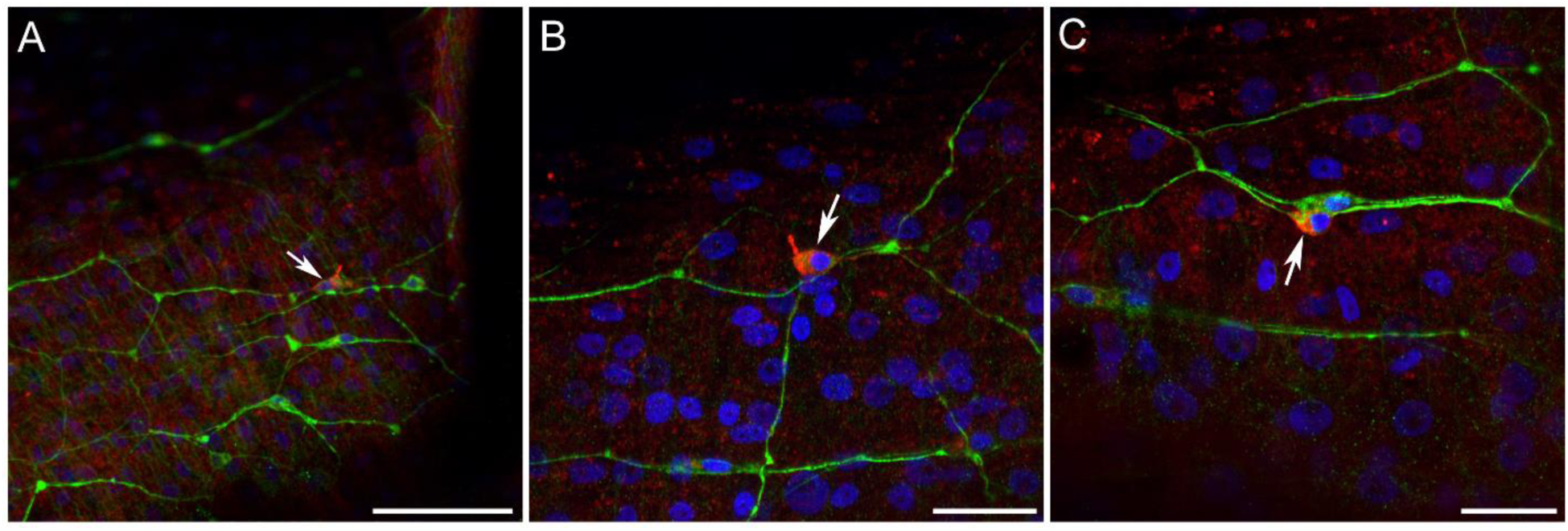
Glutamate-ir neurons in *Aequorea*. The nervous system in *Aequorea’s* subumbrella is labeled with tubulin antibody (green) and Glutamate antibody (red). DAPI is blue. A, B, C – Among the polygonal neural net stained with tubulin IR, only a small fraction of neurons is also labeled red with Glutamate IR (arrows). Because of the apical filament, we think these could be sensory-like neurons. Scale bars: A – 50 µm, B – 20 µm, C – 20 µm.

**Figure 8.**
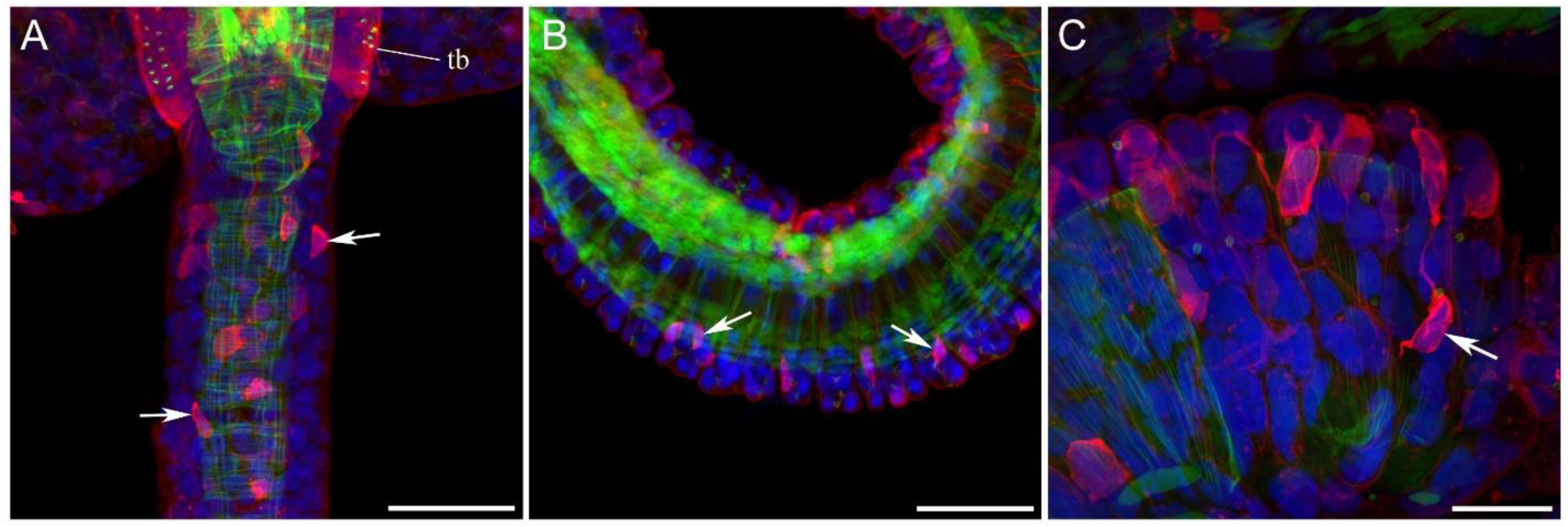
Glutamate-ir neurons in *Aglantha* tentacles. Tentacles in *Aglantha* contain many cells labeled with Glutamate IR (red), some with neuron-like morphology (have elongated processes). Phalloidin labeling is green, while DAPI is blue. A – Image of the tentacle next to the tentacle base (tb) at the subumbrella edge. Arrows point to some Glutamate-ir cells. B – Midsection of *Aglantha* tentacle. C – Higher magnification shows Glutamate-ir neural-type cells (arrow) with branching processes. Scale bars: A – 50 µm, B – 50 µm, C – 20 µm.

**Figure 9.**
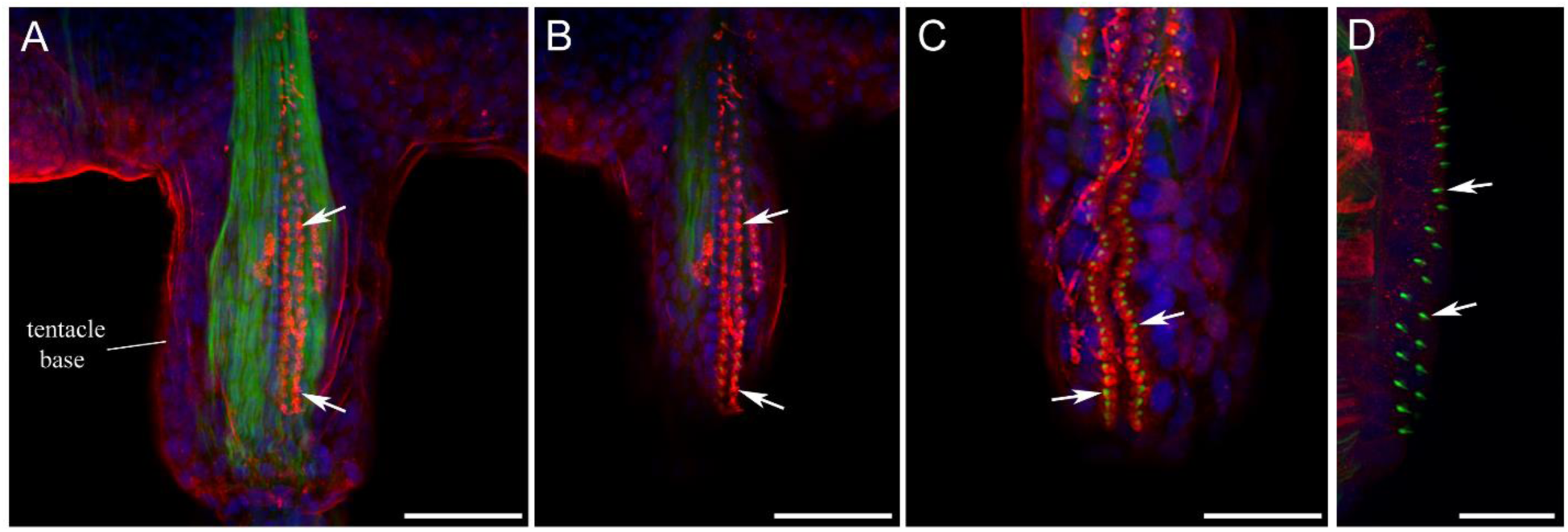
Glutamate-ir sensory cells in *Aglantha* tentacle base. Ciliated sensory cells at the base of *Aglantha* tentacles are labeled with Glutamate IR (red). Phalloidin labeling is green, while DAPI is blue. A – There are two rows of sensory cells (arrows) running longitudinally along each tentacle base. B – Optical section of the same tentacle base near the surface more clearly reveals the sensory rows. C – Higher magnification shows that each of these Glutamate-ir sensory cells has a short “thorn” in the middle labeled with phalloidin (green; shown by arrows). This is a cluster of short microvilli, which surrounds the base of long sensory cilia not visible with current labeling (see Norekian and Moroz, 2020). D – GABA antibodies do not label these sensory cells; only clusters of small microvilli are seen stained with phalloidin (green; arrows). Scale bars: A, B – 50 µm, C – 30 µm, D – 20 µm.

Glutamate IR was also observed in thin, long filaments that traverse the mesogleal region from the subumbrella swim muscle layer to the exumbrella epithelium in four species – *Aequorea, Bougainvillia, Euphysa*, and *Aglantha* (**Fig. 10**). These long Glutamate-ir filaments were not labeled with phalloidin, a universal F-actin marker (Wulf et al., 1979), and were therefore unlikely to be muscles. They may be neuronal and represent long neurites, but we cannot be certain of their identity at this point.

**Figure 10.**
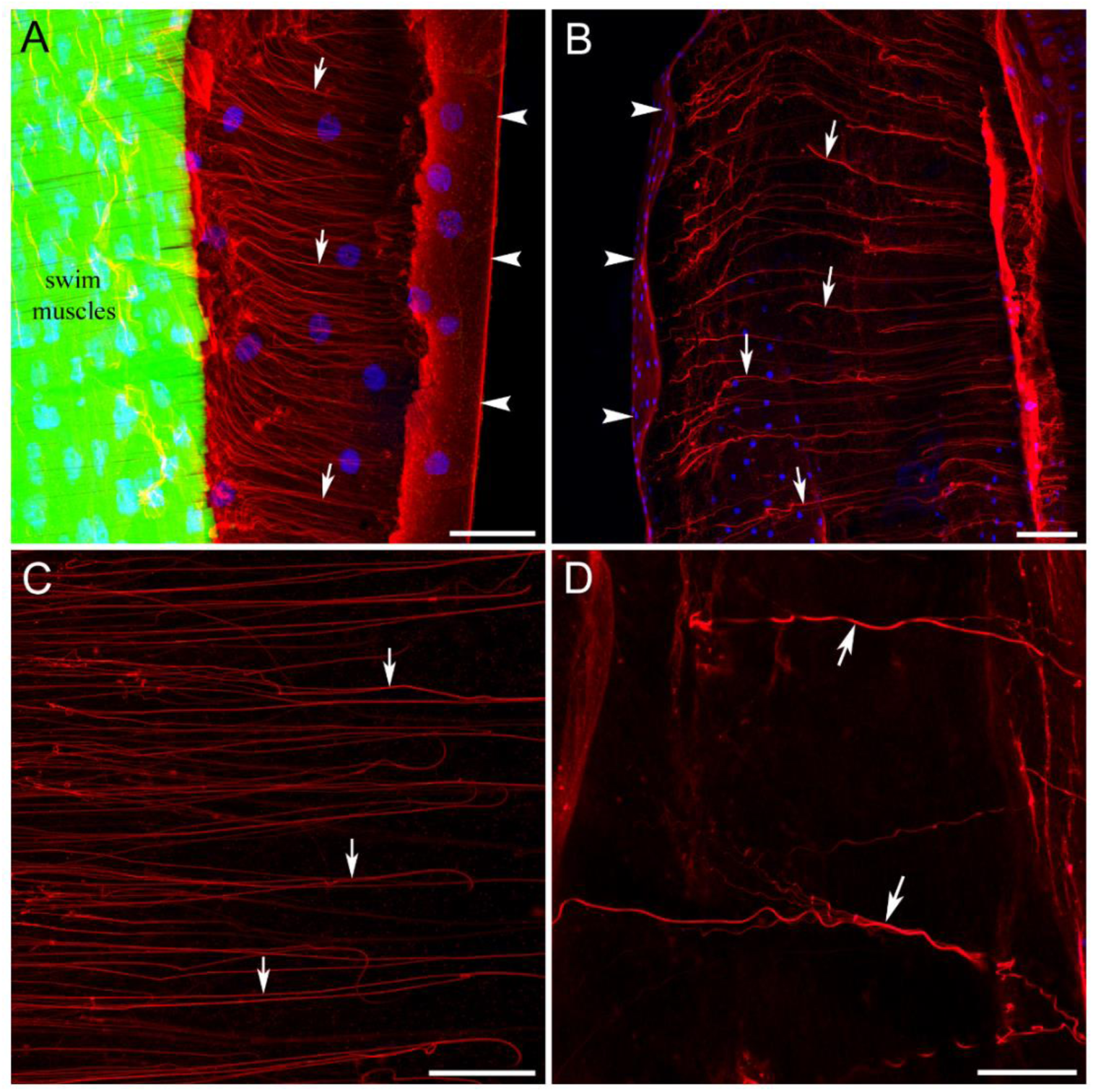
Glutamate-ir fibers in mesoglea of hydromedusae. Multiple thin fibers (arrows), which cross the umbrella wall from the exumbrella outside surface (arrowheads) to the subumbrella swim musculature, are labeled with Glutamate antibodies (red). They are not labeled with phalloidin (green) suggesting that they are probably of neural rather than muscle nature. DAPI is blue. A – Cross section of the *Aglantha* body wall. B – *Bougainvillia* body wall. C – *Euphysa* mesogleal region. D – Mesogleal region in *Aequorea*. Scale bars: A – 100 µm, B – 50 µm, C – 40 µm, D – 20 µm.

Glutamate IR also labeled specific muscle subtypes across species. For example, glutamate IR was observed in the striated muscles of *Euphysa* (Fig. 11A), the umbrella muscles of *Clytia* (Fig. 11B), some swim muscles of *Aglantha* (Fig. 11C), and the swim striated muscles of *Nanomia* nectophores (Fig. 12). GABA IR also selectively labeled muscle cells, including *Euphysa* tentacle muscles (Fig. 3B), smooth muscles along the radial canals in *Clytia* (Fig. 13A), and some striated swim muscles of *Aglantha*, along with their myoepithelial cell bodies (Fig. 13C).

**Figure 11.**
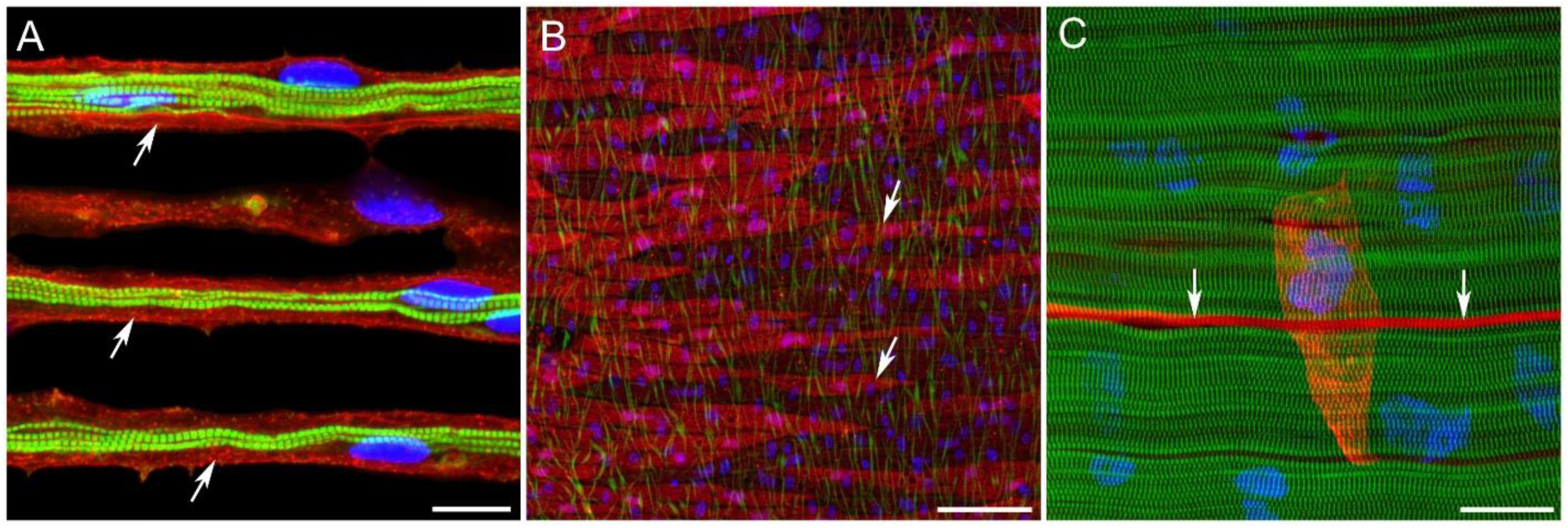
Glutamate-ir muscles in Hydrozoa. Some muscle cells are clearly labeled with Glutamate IR (red). Phalloidin labeling is green, while nuclei DAPI is blue. A – Striated muscle cells in the *Euphysa* umbrella show a strong Glutamate IR signal. B – Clusters of muscle cells in *Clytia gregaria*. C – In *Aglantha*, only a few striated swim muscle cells (arrows) show Glutamate IR. The large muscle cell body (red) that gives rise to the striated swim fibers is also visible. These cells form a continuous myoepithelial layer in subumbrella. Scale bars: A – 10 µm, B – 50 µm, C – 20 µm.

**Figure 12.**
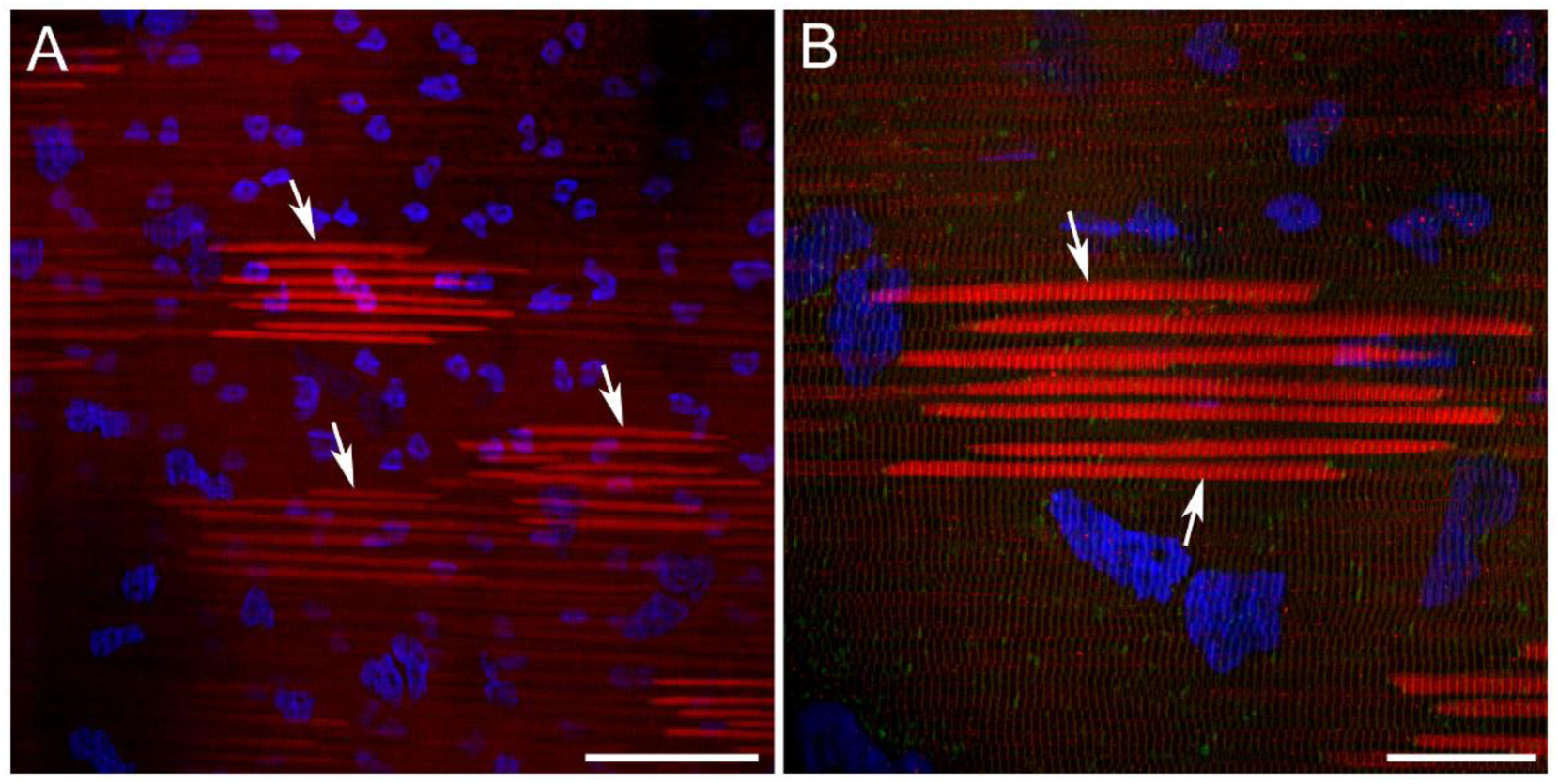
Glutamate-IR swim muscles in *Nanomia* nectophores. Glutamate IR is red; DAPI (nuclear) is blue. Some striated swim muscle fibers in *Nanomia* nectophores are Glutamate-ir. A – Different clusters of the swim muscle fibers are labeled with Glu IR at different intensities (arrows), while many other striated swim muscles do not show any Glu IR. B – Higher magnification shows a Glu-ir cluster of striated swim muscle fibers (arrows). Scale bars: A - 50 µm; B - 20 µm.

**Figure 13.**
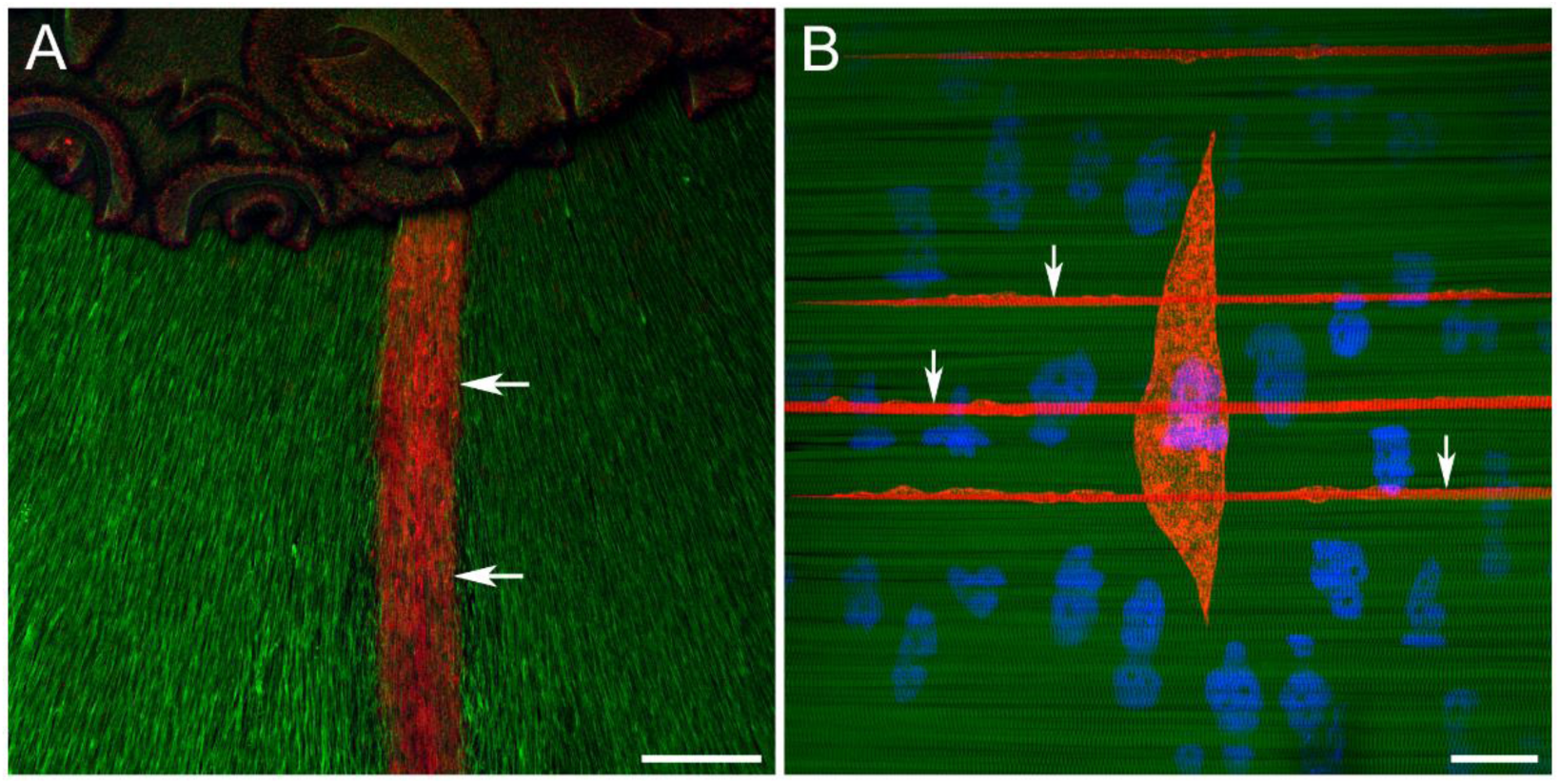
GABA IR in the muscles. GABA immunoreactivity (red) is found in some muscles across hydrozoan species. Phalloidin is green, while DAPI is blue. A – A bundle of muscles (arrows) along the radial canal in *Clytia gregaria*. The dark structure at the top is the peduncle and manubrium base. B – Striated swim muscle fibers (arrows) and their large spindle-shaped myoepithelial cell body in *Aglantha*. Scale bars: A – 200 µm, B – 20 µm.

## DISCUSSION

Despite the ambiguous phylogeny of hydrozoan medusae, the selected species reveal parallel evolution of multiple complex adaptations to pelagic life, including distinct feeding and locomotory strategies. For example, siphonophores (*Nanomia*) and trachymedusae (*Aglantha*) independently evolved rapid escape responses, with independent origins of giant axons, loss of polyp stages, and elaborate tentacle apparatuses. Representatives of the order Leptothecata (*Aequorea*, *Eutonina*, *Clytia*) are relatively slow-swimming medusae with distinct tentacles and feeding strategies. Anthoathecata medusae (*Bougainvillia*, *Polyorchis*, *Euphysa*) are distinguished by their tentacle organization and nematocyte distribution.

These complex adaptations to pelagic life in Hydrozoa may be reflected in species-specific expression patterns of glutamatergic and GABAergic cells (**Fig. 14**). Although most immunoreactive cells are not neurons, this study shows clear labeling of neuronal elements in three of the eight species studied: *Nanomia*, *Aglantha*, and *Aequorea*. Neural glutamate expression correlates with behavioral complexity. Both *Aglantha* and *Nanomia* are active swimmers with independently evolved giant axons that enable fast startle responses (Mackie, 1973; Roberts, Mackie, 1980; Mackie, 2004; Norekian, Moroz, 2020). In *Aequorea*, neural glutamate IR labeling is weak and restricted to sensory-like cells, which differ in morphology and are less abundant than in *Aglantha* and *Nanomia*. The functional role of glutamate in hydromedusae has not been investigated; however, based on morphology and localization, we can propose glutamate-dependent sensory processing in *Aglantha* and *Aequorea*, neural control of tentacles in *Aglantha*, and jet propulsion in *Nanomia*.

**Figure 14.**
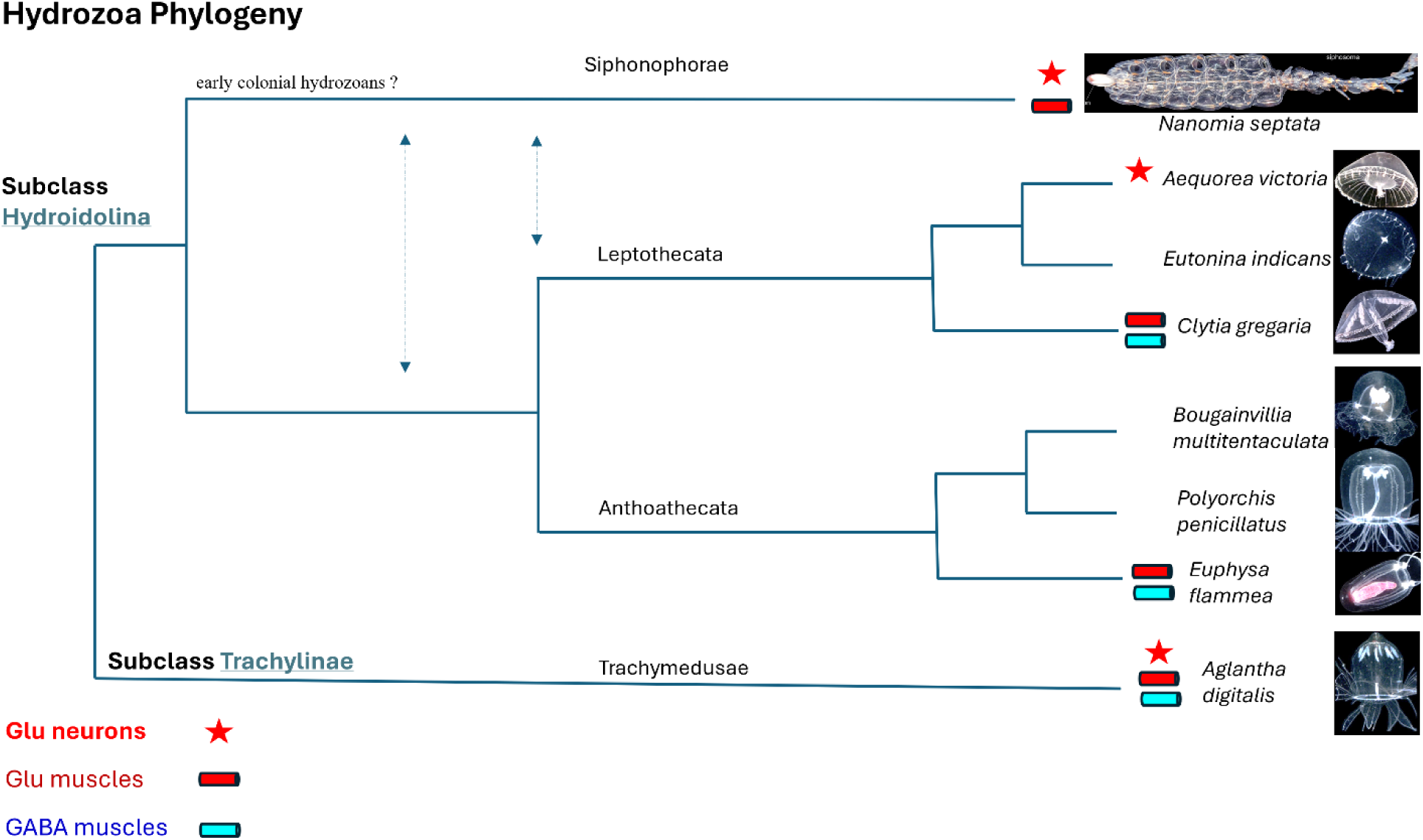
Hydrozoan phylogeny and distribution of neural and muscle elements enriched with glutamate and GABA. The schematics illustrate phylogenetic relationships of 4 orders of Hydrozoa and the presence of neurons and muscles with Glu IR and GABA IR across eighth pelagic species. The Leptothecata relationships are based on (Maronna et al., 2016); representatives of this order possess a chitinous exoskeleton or cup (theca) in their polyp stages. Anthoathecata represents species with so-called ‘naked’ polyp stages without chitinous protection. *Polyorchis* and *Bougainvillia* belong to the subclades Capileta (medusae with knobs) and Filifera (medusae without knobs), respectively. *Euphysa* belongs to the subclade Aplanulata, a lineage that lost planula stage in development. Siphonophorae are highly developed colonial hydrozoans in which individual zooids specialize into organ-like structures. Trachymedusae (*Aglantha*) secondarily lost their polyp stages, and medusae developed directly from a swimming larva. All studied medusae express distinct locomotory patterns, in terms of speed, frequency, coordination, and energetic demands. The distribution of Glu IR and GABA IR is mosaic and does not represent homologous cell types. Glu-IR neurons were found in only 3 studied species and represent different populations in terms of their locations, morphology, and perhaps functions. Glutamate IR was observed in nematocytes of all studied species. There are also several different muscle groups, which demonstrated glutamate or GABA IRs. We interpret the observed patterns in Hydrozoa as independent recruitments of glutamate and GABA in neural and muscular functions.

Across other classes of the phylum Cnidaria, putative glutamatergic neurons were observed in only one anthozoan species (*Phymactis papilosa*, Delgado et al., 2010), and non-neuronal glutamate localization was the predominant feature in the species studied (Carlyle, 1974, Anctil and Carette, 1994). Both ectodermal and endodermal distributions of putative glutamatergic cells were reported using immunolabeling and *in situ* hybridization for vesicular glutamate transporters (Oren et al., 2014; Sheloukhova, Watanabe, 2023). The sea anemone *Nematostella* has ten genes encoding vesicular glutamate transporters, whereas humans have only three and *Trichoplax* has only one (Moroz et al., 2021b). Cnidarians ‘invented’ vesicular glutamate transporters (together with placozoans) and NMDA-type receptors, including cnidaria-specific subclasses (Moroz et al., 2021). Comparative genomics also points to distinct evolutionary trajectories in the distribution of ionotropic glutamate receptors within different classes of Cnidaria; for example, the loss of akdf-type iGlu receptors in Hydrozoa (Moroz et al., 2021b).

These data, together with the highly mosaic localization patterns of glutamatergic cells, suggest that glutamate was independently recruited as a transmitter in neuronal signaling at least 3 times in Hydrozoa (**Fig. 14**), at least once in Anthozoa, and perhaps in other cnidarian classes (to be investigated). Ctenophores also appear to have independently recruited glutamate as a neuromuscular transmitter (Moroz et al., 2014; Moroz, Norekian, 2026). Conservatively, combining these data with those on Bilaterians, we might recognize at least 6 origins of glutamatergic neurons.

This broad evolutionary plasticity of glutamatergic signaling may be explained by two factors. First, it is relatively easy to “make” glutamatergic cells - this process requires expression of a gene encoding a vesicular glutamate transporter, as shown by Takamori et al. (2000). Such ‘simplicity’ may underlie the observed versatility and diversification of glutamatergic systems, as we discover in Hydrozoa. This is a hypothesis about “*how*” glutamatergic systems might have arisen in cnidarian neurons and elsewhere. But *why* was glutamate recruited into the secretory machinery of neurons, and other cell types? The second factor may be related to glutamate’s role as an anaplerotic substrate that refills the TCA cycle, in which the uptake and oxidation of one molecule of glutamate yield 23-26 molecules of ATP (McKenna, 2013).

These two aspects of glutamate biochemistry, together with its roles as an osmolyte and chaperone, might contribute to its preferential selection in muscular and neural systems, especially in species and cells under energetic constraints across various ecological niches, as in *Nanomia* and *Aglantha*. Predictably, intense contractions of striated swim muscles in these species are energy-demanding (as in human muscles at the onset of exercise) and require anaplerotic replenishment of the catalytic intermediates of the TCA cycle (Brunengraber, Roe, 2006). Similar metabolic functions of glutamate have been widely reported in neuron-glia interplay (McKenna, 2013; Andersen, 2025).

The shared developmental trajectories of neurons and nematocytes are of considerable evolutionary and physiological importance (Babonis et al., 2022; Steger et al., 2022; Babonis, 2025). Nematocytes contain an extraordinarily high concentration of poly-gamma-glutamate, reaching 2 M in *Hydra* (Weber, 1990). Notably, cnidarians directly acquire the poly-glutamate synthase gene from bacteria via horizontal gene transfer (Denker et al., 2008). Exogenous Glu is also reported to increase stenotele discharge in *Hydra* (Scappaticci et al., 2010; Scappaticci and Kass-Simon, 2008), but positive feedback from Glu released during cnidocyte discharge might also contribute to coordinated feeding/defensive behaviors (Lenhoff and Bovaird, 1961; Kass-Simon et al., 2003, Kay, Kass-Simon, 2009; Pierobon, 2012; Pierobon et al., 2004).

During development, the vesicular glutamate transporter was expressed in two distinct populations of *Hydra* nematoblasts (Moroz et al., 2021), where glutamate uptake may contribute to polymer synthesis (i.e., poly-gamma-glutamate). Single-cell transcriptomics suggest that only a small fraction of putative ectodermal neural or myoepithelial cells might contain vesicular glutamate transporters and be glutamatergic.

The distribution and functions of GABA in hydrozoans remain enigmatic; its coupling to glutamate metabolism, particularly in muscles, warrants systematic study. In addition, we observed other putative glutamatergic and GABAergic cell types (**Figs. 3, 5, 10**) with unknown physiology. Here, we also anticipate distinct metabolic cell states under transient conditions of elevated glutamate or GABA synthesis.

### Conclusion, Limitations and Future Directions

Comparative data clearly indicate selective recruitment of glutamate and GABA in neural and muscular functions among cnidarians. Given the need to study the physiological roles of glutamate and GABA in behavioral integration, a major bottleneck is detecting vesicular or non-vesicular *release* of these molecules from specific cell populations (e.g., glutamate release following nematocyte discharge, muscle contractility, and neural physiological activation).

Beyond its transmitter roles, glutamate functions as an osmolyte and chaperone for protein complexes and protein-DNA interactions (Deredge et al., 2010; Young, Ajami, 2000), and polyglutamylation is functionally essential (Mahalingan et al., 2020; Martinez-Limon et al., 2016; Regnard et al., 2000). These chemical modifications might be critical for the origin of nematocytes, in particular, and for neuronal origins in general.

We strongly advocate a broad comparative survey of glutamate and GABA signaling across all eight cnidarian classes, leveraging their remarkable diversity of solutions for coping with complex environmental perturbations. This will not only advance evolutionary reconstructions but also yield discoveries of novel molecular and system functions that inform synthetic biology.

## Acknowledgments

We thank FHL for their excellent facilities, including the Nikon Laser Scanning confocal microscope. This research was supported by the National Science Foundation grant (IOS-2341882) and National Institute of Health grant (5R01NS11449) to LLM.

## Author contributions

Study design, conceptualization and funding (LLM), immunohistochemical staining (TPN), data analysis and writing (LLM, TPN).

## Data Availability Statement

The data that support the findings of this study are available from the corresponding author upon reasonable request.

## REFERENCES

Andersen, J.V., 2025. The Glutamate/GABA-Glutamine cycle: Insights, updates, and advances. J Neurochem. 169(3):e70029. doi: 10.1111/jnc.70029.

Anctil M. 2009. Chemical transmission in the sea anemone *Nematostella vectensis*: A genomic perspective. Comp Biochem Physiol Part D Genomics Proteomics. 4(4):268–289. doi: 10.1016/j.cbd.2009.07.001.

Anctil, M., 2015. Dawn of the Neuron: The Early Struggles to Trace the Origin of Nervous Systems, Montreal: McGill-Queen’s University Press. ISBN: 9780773597327. doi: 10.1515/9780773597327

Anctil, M., Carette, J. P., 1994. Glutamate immunoreactivity in non-neuronal cells of the sea anemone *Metridium senile*. Biol Bull 187, 48–54.

Babonis, L.S., 2025. That stinging sensation: Modularity and the origin of the stinging cell. Integr Comp Biol. Sep 26;65(3):661-675. doi: 10.1093/icb/icaf070.

Babonis, L.S., Enjolras, C., Ryan, J.F., Martindale, M.Q., 2022. A novel regulatory gene promotes novel cell fate by suppressing ancestral fate in the sea anemone *Nematostella vectensis*, Proc. Natl. Acad. Sci. U.S.A. 119 (19) e2113701119. Doi: 10.1073/pnas.2113701119.

Bosch, T.C.G., Klimovich, A., Domazet-Loso, T., Grunder, S., Holstein, T.W., Jekely, G., Miller, D.J., Murillo-Rincon, A.P., Rentzsch, F., Richards, G.S., Schroder, K., Technau, U., Yuste, R., 2017. Back to the Basics: Cnidarians Start to Fire. Trends Neurosci 40, 92–105.

Brunengraber, H., Roe, C. R., 2006. Anaplerotic molecules: current and future. J Inherit Metab Dis 29, 327–331.

Carlyle, R. F., 1974. The occurrence in and actions of amino acids on isolated supra oral sphincter preparations of the sea anemone *Actinia equina*. J Physiol 236, 635–652.

Delgado, L. M., Couve, E., Schmachtenberg, O., 2010. GABA and glutamate immunoreactivity in tentacles of the sea anemone *Phymactis papillosa* (LESSON 1830). J Morphology 271, 845–852.

Denker, E., Bapteste, E., Le Guyader, H., Manuel, M., Rabet, N., 2008. Horizontal gene transfer and the evolution of cnidarian stinging cells. Curr Biol 18, R858–859.

Deredge, D. J., Baker, J. T., Datta, K., Licata, V. J., 2010. The glutamate effect on DNA binding by pol I DNA polymerases: osmotic stress and the effective reversal of salt linkage. J Mol Biol 401, 223–238.

Kass-Simon, G., Pannaccione, A., Pierobon, P., 2003. GABA and glutamate receptors are involved in modulating pacemaker activity in *Hydra*. Comparative Biochemistry and Physiology Part A: Molecular & Integrative Physiology 136, 329–342.

Kelava, I., Rentzsch, F., Technau, U., 2015. Evolution of eumetazoan nervous systems: insights from cnidarians. Phil. Trans. R. Soc. B 19 December 2015; 370 (1684): 20150065. 10.1098/rstb.2015.0065

Lenhoff, H., Bovaird, J., 1961. Action of glutamic acid and glutathione analogues on the *Hydra* glutathione-receptor. Nature 189, 486–487.

Kay, J. C., Kass-Simon, G., 2009. Glutamatergic transmission in hydra: NMDA/D-serine affects the electrical activity of the body and tentacles of *Hydra vulgaris* (Cnidaria, Hydrozoa). Biol Bull 216, 113–125

Mackie, G.O., 1970. Neuroid conduction and the evolution of conducting tissues. Q Rev Biol. 45(4):319–32. doi: 10.1086/406645.

Mackie, G.O., 1973. Report on giant nerve fibres in *Nanomia*. Publications of the Seto Marine Biological Laboratory; 20:745–756.

Mackie, G. O., 1990. The elementary nervous systems revisited. American Zoologist 30, 907–920.

Mackie, G.O. 2004. Central neural circuitry in the jellyfish *Aglantha*: a model ‘simple nervous system’. Neurosignals. 13(1-2):5–19. doi: 10.1159/000076155.

Mahalingan, K. K., Keith Keenan, E., Strickland, M., Li, Y., Liu, Y., Ball, H. L., Tanner, M. E., Tjandra, N., Roll-Mecak, A., 2020. Structural basis for polyglutamate chain initiation and elongation by TTLL family enzymes. Nat Struct Mol Biol 27, 802–813.

Marlow, H.Q., Srivastava, M., Matus, D.Q., Rokhsar, D., Martindale, M.Q., 2009. Anatomy and development of the nervous system of *Nematostella vectensis*, an anthozoan cnidarian. Developmental Neurology 69 (4), 235–254. doi: 10.1002/dneu.20698

Maronna, M., Miranda, T., Peña Cantero, Á., Barbeitos, M.S., Marques, A.S., 2016. Towards a phylogenetic classification of Leptothecata (Cnidaria, Hydrozoa). Sci Rep 6, 18075. doi: 10.1038/srep18075

Martinez-Limon, A., Alriquet, M., Lang, W. H., Calloni, G., Wittig, I., Vabulas, R. M., 2016. Recognition of enzymes lacking bound cofactor by protein quality control. Proc Natl Acad Sci U S A 113, 12156–12161.

McKenna, M. C., 2013. Glutamate pays its own way in astrocytes. Front Endocrinol (Lausanne) 4, 191.

McLaggan, D., Naprstek, J., Buurman, E. T., Epstein, W., 1994. Interdependence of K^+^ and glutamate accumulation during osmotic adaptation of *Escherichia coli*. J Biol Chem 269, 1911–1917.

Moroz, L.L. 2014. The genealogy of genealogy of neurons. Commun Integr Biol 7(6):e993269. doi: 10.4161/19420889.2014.993269.

Moroz, L.L. 2021. Multiple origins of neurons from secretory cells. Front Cell Dev Biol; 9:669087; doi: 10.3389/fcell.2021.669087.

Moroz, L. L., Kocot, K. M., Citarella, M. R., Dosung, S., Norekian, T. P., Povolotskaya, I. S., Grigorenko, A. P., Dailey, C., Berezikov, E., Buckley, K. M., Ptitsyn, A., Reshetov, D., Mukherjee, K., Moroz, T. P., Bobkova, Y., Yu, F., Kapitonov, V. V., Jurka, J., Bobkov, Y. V., Swore, J. J., Girardo, D. O., Fodor, A., Gusev, F., Sanford, R., Bruders, R., Kittler, E., Mills, C. E., Rast, J. P., Derelle, R., Solovyev, V. V., Kondrashov, F. A., Swalla, B. J., Sweedler, J. V., Rogaev, E. I., Halanych, K. M., Kohn, A. B. 2014. The ctenophore genome and the evolutionary origins of neural systems. Nature 510, 109–114. doi: 10.1038/nature13400

Moroz, L.L., Kohn, A.B. 2015. Unbiased view of synaptic and neuronal gene complement in ctenophores: Are there pan-neuronal and pan-synaptic genes across Metazoa? Integr Comp Biol 55, 1028–1049. doi: 10.1093/icb/icv104.

Moroz, L. L., Kohn, A. B., 2016. Independent origins of neurons and synapses: insights from ctenophores. Philos Trans R Soc Lond B Biol Sci 371, 20150041. doi: 10.1098/rstb.2015.0041.

Moroz, L.L., Meech, R.W., Sweedler, J.V., Mackie, G.O., 2004. Nitric oxide regulates swimming in the jellyfish *Aglantha digitale*. Journal of Comparative Neurology 471 (1), 26–36. doi: 10.1002/cne.20023

Moroz, L.L., Nikitin, N.A., Poličar, P.G., Andrea B. Kohn, A.B., Romanova, D.Y. 2021a. Evolution of glutamatergic signaling and synapses. Neuropharmacology;199:108740. doi: 10.1016/j.neuropharm.2021.108740.

Moroz, L.L., Norekian, T.P., 2026. Glutamatergic systems in ctenophores. bioRxiv, 2026.08. 09.743775. doi: 10.64898/2026.08.09.743775

Moroz, L. L., Romanova, D. Y., Kohn, A. B. 2021b. Neural versus alternative integrative systems: molecular insights into origins of neurotransmitters. Philos Trans R Soc Lond B Biol Sci 376, 20190762. doi: 10.1098/rstb.2019.0762.

Moroz, L. L., Sohn, D., Romanova, D. Y., Kohn, A. B., 2020. Microchemical identification of enantiomers in early-branching animals: Lineage-specific diversification in the usage of D-glutamate and D-aspartate. Biochem Biophys Res Commun 527, 947–952. doi: 10.1016/j.bbrc.2020.04.135.

Norekian, T.P., Moroz, L.L., 2020. Atlas of the neuromuscular system in the Trachymedusa *Aglantha digitale*: Insights from the advanced hydrozoan. J Comp Neurol. 528(7):1231–1254. doi: 10.1002/cne.24821.

Norekian TP, Moroz, L.L. 2024. Recording cilia activity in ctenophores. Methods Mol Biol. 2024;2757:307–313. doi: 10.1007/978-1-0716-3642-8_14.

Norekian, T.P., Moroz, L.L., 2026. Neuromuscular architecture of the siphonophore colony. bioRxiv, 2026.08. 03.742546. 10.64898/2026.08.03.742546

Oren, M., Brickner, I., Appelbaum, L., Levy, O., 2014. Fast neurotransmission related genes are expressed in non nervous endoderm in the sea anemone *Nematostella vectensis*. PLoS One 9, e93832.

Pierobon, P., 2012. Coordinated modulation of cellular signaling through ligand-gated ion channels in *Hydra vulgaris* (Cnidaria, Hydrozoa). Int J Dev Biol 56, 551–565.

Pierobon, P., Sogliano, C., Minei, R., Tino, A., Porcu, P., Marino, G., Tortiglione, C., Concas, A., 2004. Putative NMDA receptors in *Hydra*: a biochemical and functional study. Eur J Neurosci 20, 2598–2604.

Regnard, C., Desbruyeres, E., Huet, J. C., Beauvallet, C., Pernollet, J. C., Edde, B., 2000. Polyglutamylation of nucleosome assembly proteins. J Biol Chem 275, 15969–15976.

Roberts, A., Mackie, G.O., 1980. The giant axon escape system of a hydrozoan medusa, *Aglantha digitale*. J Exp Biol. 84:303–18. doi: 10.1242/jeb.84.1.303.

Scappaticci, A. A., Jr., Kahn, F., Kass-Simon, G., 2010. Nematocyst discharge in *Hydra vulgaris*: Differential responses of desmonemes and stenoteles to mechanical and chemical stimulation. Comp Biochem Physiol A Mol Integr Physiol 157, 184–191.

Scappaticci, A. A., Kass-Simon, G., 2008. NMDA and GABA B receptors are involved in controlling nematocyst discharge in *Hydra*. Comp Biochem Physiol A Mol Integr Physiol 150, 415–422.

Sheloukhova, L., Watanabe, H., 2023. Analysis of cnidarian Gcm suggests a neuronal origin of glial EAAT1 function. Sci Rep. 13(1):14790. doi: 10.1038/s41598-023-42046-9.

Steger, J., Colem, A.G., Denner, A., Lebedeva, T., Genikhovich, G., Ries, A., Reischl, R., Taudes, E., Lassnig, M., Technau, U., 2022. Single-cell transcriptomics identifies conserved regulators of neuroglandular lineages. Cell Rep. 40(12):111370. doi: 10.1016/j.celrep.2022.111370.

Takamori, S., Rhee, J. S., Rosenmund, C., Jahn, R., 2000. Identification of a vesicular glutamate transporter that defines a glutamatergic phenotype in neurons. Nature 407, 189–194.

Young, V. R., Ajami, A. M., 2000. Glutamate: an amino acid of particular distinction. J Nutr 130, 892S–900S.

Wang, W., Jack, B. M., Wang, H. H., Kavanaugh, M. A., Maser, R. L., Tran, P. V., 2021. Intraflagellar transport proteins as regulators of primary cilia length. Frontiers in Cell and Developmental Biology 9.

Weber, J., 1990. Poly(gamma-glutamic acid)s are the major constituents of nematocysts in *Hydra* (Hydrozoa, Cnidaria). J Biol Chem 265, 9664–9669.

Wehland J, Willingham MC, Sandoval IV. 1983. A rat monoclonal antibody reacting specifically with the tyrosylated form of alpha-tubulin. I. Biochemical characterization, effects on microtubule polymerization in vitro, and microtubule polymerization and organization in vivo. J Cell Biol. Nov;97(5 Pt 1):1467-75. doi: 10.1083/jcb.97.5.1467.

Wehland J, Willingham MC. 1983. A rat monoclonal antibody reacting specifically with the tyrosylated form of alpha-tubulin. II. Effects on cell movement, organization of microtubules, and intermediate filaments, and arrangement of Golgi elements. J Cell Biol. Nov;97(5 Pt 1):1476-90. doi: 10.1083/jcb.97.5.1476.

Wulf E, Deboben A, Bautz FA, Faulstich H, Wieland T. 1979. Fluorescent phallotoxin, a tool for the visualization of cellular actin. Proc Natl Acad Sci U S A. Sep;76(9):4498–502. doi: 10.1073/pnas.76.9.4498.

